# Gene network centrality affects parallel evolution and local adaptation in wild yeast

**DOI:** 10.64898/2026.03.11.711091

**Authors:** Swapna K. Subramanian, Daniel I. Bolnick

**Affiliations:** University of Connecticut, Department of Ecology and Evolutionary Biology, Storrs, CT, 06269, USA

**Keywords:** parallel evolution, wild yeast, evolutionary genomics, population genetics, gene-gene interaction networks

## Abstract

The predictability of evolution remains a fundamental question in biology. While parallel evolution has been observed across taxa, we lack a mechanistic framework for predicting when evolution will be repeatable versus contingent. Here, we test whether gene network architecture predicts evolutionary repeatability and local (mal)adaptation in natural populations. We sequenced 38 wild *Hanseniaspora uvarum* yeast strains from two replicated apple varieties across four Connecticut orchards, yielding 29,489 high-quality SNPs. Population genomic analyses suggested that genetic differentiation was structured primarily by orchard environment. Reciprocal transplant experiments across all orchard-variety combinations demonstrated local (mal)adaptation at multiple ecological scales, with strongest (mal)adaptive responses at the orchard-variety level. By mapping *H. uvarum* genes to the comprehensive *Saccharomyces cerevisiae* genetic interaction network, we found that network centrality affects evolutionary outcomes. Highly connected genes occupying bow-tie network positions evolved in parallel across replicated apple varieties and central genes harbored alleles with adaptive fitness effects. In contrast, peripheral genes with fewer interactions facilitated rapid, non-parallel evolution to orchard-variety interactions and were associated with local maladaptation. Analysis of 150-gene sets evolving to apple variety within each orchard revealed greater-than-random similarity at genetic, functional, and network levels, with network-level predictability significantly exceeding gene-level predictability. These results suggest that gene network architecture could provide a mechanistic foundation for determining which evolutionary changes will be repeatable, potentially enabling evolutionary predictions even when specific genes cannot be identified.

## Introduction

The predictability of evolution remains one of biology’s most fundamental questions. Recent advances suggest that evolutionary repeatability exists at different biological scales (genotypic, phenotypic, pathway, etc.), but is inconsistent (Bolnick et al. 2018, Pearless and Freed 2024). This raises the question of why evolution is sometimes repeatable, and sometimes not? (Bolnick et al. 2018). Many variables impact our ability to predict evolution, including mutation rate, population size, ecology, strength of selection, and eco-evolutionary dynamics (Pearless and Freed 2024). Historical contingency has also been recognized as a potential constraint on evolution since Wright’s adaptive landscape (Wright 1932). Subsequent work on hybrid incompatibility, Gould’s “tape of life”, and Lenski’s LTEE expanded this framework (Gould 1989, Fierst and Hanen 2010, Blount et al. 2018, Pearless and Freed 2024). This body of work points to the significance of genetic interactions, or epistasis, on evolutionary potential and where in the genome evolution is most likely to occur. These works raise the question of whether epistasis could serve as a possible explanation for why adaptation sometimes occurs in parallel and sometimes does not.

Epistatic interactions are theorized to influence where evolution occurs in the genome, its rate, and its repeatability (Yang and Scarpino 2020). Genes involved in essential processes tend to be more connected in gene-gene interaction networks and have more negative epistatic interactions that limit accessible mutational paths (Costanzo et al. 2016). This connectivity results in slower evolution due to greater chance of deleterious effects of mutations (Fraser et al. 2002, Alvarez-Ponce et al. 2017), it also means that central genes are more likely to evolve in parallel. More specifically, central genes occupying bridging positions and facilitating information exchange, also known as “bow-tie” genes, are thought to be “evolutionary hot-spots” for repeatability (Friedlander et al. 2015). Bow-tie genes have been empirically shown to evolve repeatably across *Drosophila* species (Stern and Orgogozo 2009, Friedlander et al. 2015). In contrast, genes with fewer gene-gene interactions could evolve faster, responding to imminent selection pressures due to their potential to harbor more mutations and reduced negative epistatic effects. These peripheral genes could therefore be more likely to evolve in non-parallel, lineage-specific ways in response to localized selection pressures. Epistasis is also theorized to affect which genes are involved in local adaptation versus maladaptation.

The locations in which mutations occur and the fitness effects of those mutations has been shown to be influenced by epistasis, but there is conflicting evidence as to what effect network position has on adaptive value (Olson-Manning et al. 2013). Central genes are thought to evolve in response to more predictable, consistent selection pressures, likely resulting in adaptive outcomes, especially since deleterious alleles in central genes are far more likely to be pruned out (Kimura and Ohta 1974). In contrast, peripheral genes are thought to respond more to ephemeral, rapidly changing selection pressures. Therefore, there is more potential for peripheral genes to evolve in directions that may not be adaptive long-term, resulting in maladaptation (Brady et al. 2019). Theoretical models suggest gene centrality should impact evolutionary predictability (Des Marais et al. 2017, Yang and Scarpino 2020). Empirical examples in gene regulatory networks and protein interaction networks generally support these model predictions (Jovelin and Phillips 2009, Alvarez-Ponce 2017). However, these hypotheses remain difficult to test empirically in natural systems due to two key limitations: 1) few natural systems provide robust parallel and non-parallel selection pressures in replicated contexts, and 2) extremely limited information on gene-gene interactions for most organisms. Here, we present evidence of evolution driven by network centrality by evaluating wild yeast in apple orchards from clonal apple varieties and using the already established yeast gene-gene interaction network (Costanzo et al. 2016).

Wild yeast populations in Connecticut apple orchards provide an ideal “natural” experimental system to test whether gene network centrality impacts evolutionary repeatability and adaptation. This system combines the advantages of meticulously maintained generational orchards with the evolutionary complexity of natural populations. Apple orchards create variable environmental treatments for the replicated habitats of clonal apple varieties growing within them. Wild yeast colonizing these apples have the potential to be evolving to these different scales of selection pressures. With the same apple varieties growing in different orchard treatments, we can measure parallel evolution to apple variety across orchards, local evolution to each orchard environment, and condition-specific non-parallel evolution driven by orchard-variety interactions. Wild yeast genomic studies have revealed that vineyard-associated wild yeasts exhibit strong geographic clustering and local adaptation patterns (Gayevskiy and Goddard 2012, Albertin et al. 2016). Yeast are also widely used to conduct laboratory evolution experiments regarding repeatability of evolution (Segre et al. 2006, Lang et al. 2013) and have been demonstrated to evolve rapidly to selection pressures. Yeast evolution is sometimes parallel and sometimes not, with variation in parallelism across biological scales (Dhar et al. 2012, Gerstein et al. 2012).

A huge limitation for understanding the effect of epistasis on evolutionary outcomes and repeatability has been the difficulty in determining the epistatic relationships of genes in most organisms. This is another advantage of studying yeast, as there is not only extensive genetic information available for yeast, but the development of the comprehensive yeast gene-gene interaction network (Costanzo et al. 2016) has provided information on how all genes in the yeast genome interact with each other. These advancements make studying wild yeast in apple orchards an ideal system for evaluating the effect of gene-gene interactions on the predictability of evolution in natural systems.

Here, we leveraged wild *Hanseniaspora uvarum* yeast evolving across Connecticut apple orchards to test whether gene network position predicts evolutionary outcomes. We hypothesized that gene network architecture determines when evolution will be repeatable versus contingent in natural populations. Specifically, we predicted that bow-tie genes would evolve in parallel across replicated apple varieties, and less connected genes would evolve in non-parallel to orchard-variety interaction. We further predicted that parallele evolution in central genes would be associated with beneficial adaptive effects, while peripheral genes would harbor more negative maladaptive alleles. We found that highly connected genes occupying bow-tie network positions evolve in parallel across apple varieties and are involved in adaptive fitness effects. We additionally found that peripheral genes with fewer interactions facilitate non-parallel evolution and are associated with local maladaptation. These results suggest that network architecture provides a mechanistic foundation for forecasting which evolutionary changes will be repeatable versus contingent in natural populations.

## Results and Discussion

### *H. uvarum* yeast evolved genetic differences among orchards and apple varieties

We sequenced 38 *H. uvarum* strains from eight populations (four orchards x two apple varieties) using PacBio HiFi technology, yielding 29,489 high-quality SNPs after linkage disequilibrium pruning (SI Appendix, Fig. S1). Each population represents 3-5 yeast strains from a specific orchard-variety combination. We analyzed population structure using discriminant analysis of principal components (DAPC), which suggested that genomic differentiation is structured mostly by orchard differences. Among orchards, the first discriminant axis captured 79.0% of the genetic variation that distinguishes strains, with the second axis capturing 17.7% (Fig. 1A-B). A discriminant analysis to distinguish varieties (ignoring orchard) yields a single axis, with overlapping distributions between Cortland and Golden Delicious strains (Fig. 1A-B). PERMANOVA confirmed significant differentiation by orchard (*F* = 2.30, *p* = 0.001, *R²* = 0.169) but not by apple variety (*F* = 1.17, *p* = 0.24, *R²* = 0.031).

**Figure. 1.**
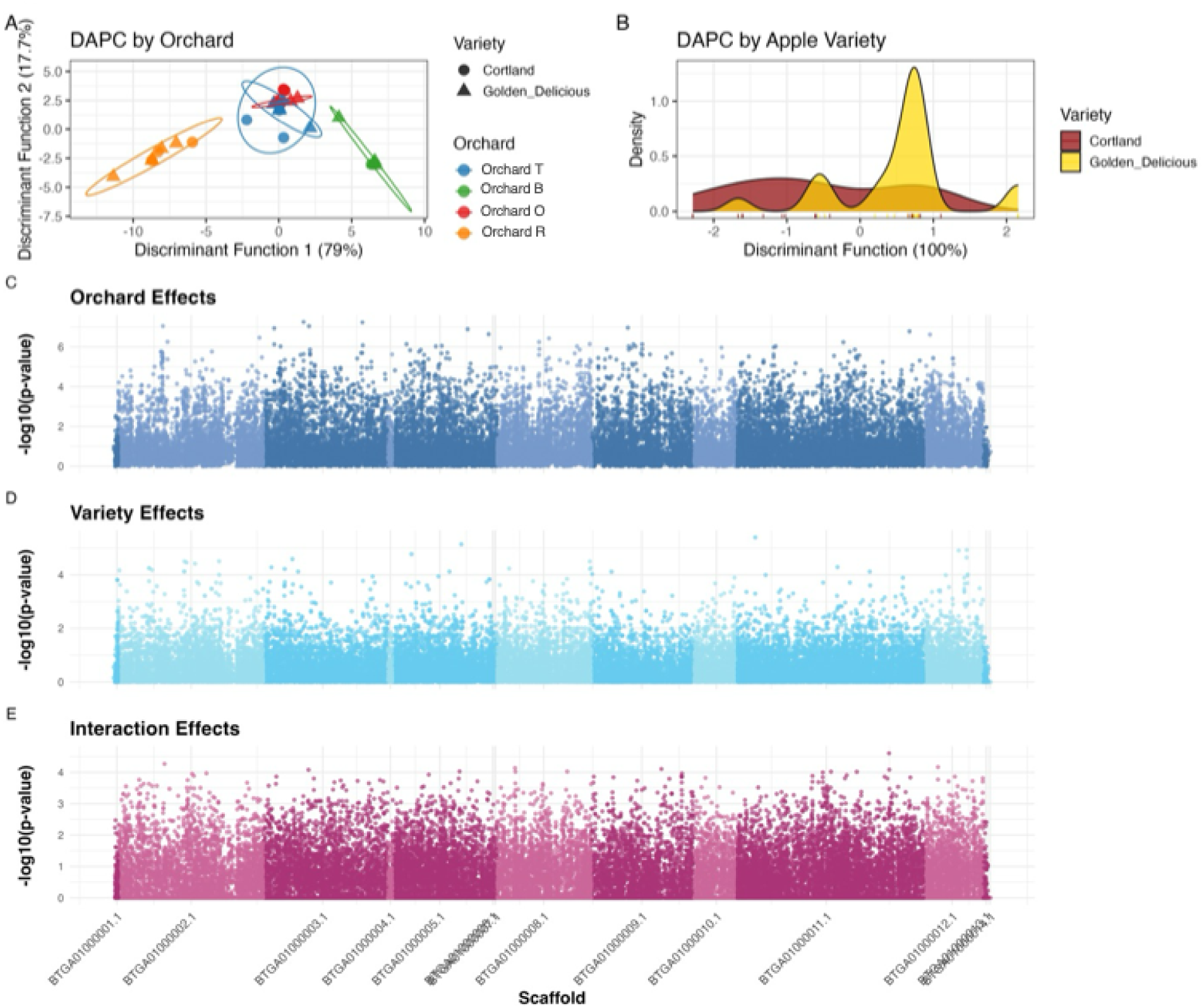
Population structure and genomic landscape of *Hanseniaspora uvarum* across Connecticut apple orchards. (A) Discriminant Analysis of Principal Components (DAPC) reveals genetic differentiation by orchard (left) and apple variety (right). Orchard DAPC explains 79% and 17.7% of variance on the first two discriminant functions, while variety DAPC explains 100% of variance on a single discriminant function. (B-D) Manhattan plots show genome-wide association patterns across 14 scaffolds for orchard effects (B), variety effects (C), and orchard×variety interaction effects (D).

To quantify the magnitude of genetic differentiation, we calculated F_ST_ at orchard, variety, and orchard-variety scales, and used binomial generalized linear models to determine which factors explain genomic variation across individual SNPs. For each SNP, we calculated variance explained as the proportional deviance reduction for each factor relative to the total model deviance. Orchard-variety differentiation patterns showed mean F_ST_ = 0.06 across all 8 orchard-variety combinations. When grouping by orchard (collapsing across varieties within orchards), F_ST_ = 0.036; when grouping by variety (collapsing across orchards within varieties), F_ST_ = 0.006. The higher total differentiation across all 8 orchard-variety combinations reflects the combined effects of orchard, variety, and orchard×variety factors, indicating stronger differentiation among orchards than between apple varieties. Genome-wide association (GWAS) analyses of SNPs revealed that 31.9% of SNPs showed orchard associations, 11.3% showed apple variety associations, and 21.2% exhibited significant interactions between orchard and variety (*p* < 0.05) (Fig 1C-E;, SI Appendix, Fig. S2). Proportion tests confirmed that all three categories significantly exceeded the 5% expectation by false discovery alone (all *p* < 2e-16), indicating genuine genomic differentiation rather than statistical noise (SI Appendix, Fig. S2, top genes in Table S1).

Variance partitioning also confirmed that genomic variation is primarily structured by orchard-level factors (52.0% mean variance explained), with variety contributing 13.7% and orchard × variety interactions accounting for 34.4% of remaining variation (Fig 1C-E).

These patterns suggest that orchard-level genetic differentiation could reflect stronger selection pressures from orchard environments (climate, soil, management practices) compared to apple variety biochemistry. More likely, substantial gene flow between apple varieties (which are often meters apart within a given orchard) compared to orchards (which are separated by tens of kilometers) reduces variety level differentiation. Orchard-level factors such as geographic isolation, distinct management practices (including fungicide identity and application schedules), and microclimate differences can create strong barriers to gene flow, leading to pronounced orchard-variety combination structure (Slatkin 1987). The substantial orchard variety interactions could indicate context-dependent evolutionary responses, where each orchard-variety combination creates unique selective environments that cannot be predicted from either factor alone. The interactions could also indicate historical contingency, as isolated orchards may have different genotypes available, constraining how yeast can adapt to a given apple variety (Blount et al. 2018).

Low differentiation between apple varieties could reflect several processes at work. Gene flow may homogenize allele frequencies between varieties within orchards more effectively than between orchards (Slatkin 1987, Whitlock and McCauley 1999, Lenormand 2002), as yeast dispersal by insects, wind, or farming equipment likely occurs more frequently within than between orchards. Alternatively, apple varieties could have relatively similar selective environments despite differences in their biochemical composition. While apple cultivars differ in sugar profiles, organic acids, and secondary metabolite concentrations (Wu et al. 2007, Aprea et al. 2017), these biochemical differences may exert weaker selective pressures on yeast orchard-variety combinations than orchard level environmental pressures. Variety-level differentiation could also be masked by non-parallel evolutionary responses to variety-specific selection pressures, where different genetic variants orchards produce similar adaptive outcomes to the same apple-variety characteristics.

### Identifying loci associated with local (mal) adaptation to orchards and apple varieties

We tested whether genetic differentiation reflects adaptive evolution by testing fitness effects of moving yeast between apples in a reciprocal transplant experiment. All 40 strains were grown in full-factorial design on fresh apples from all eight orchard-variety combinations (SI Appendix, Supplemental Fig. S3).

We measured yeast performance as the diameter of decayed apple tissue after seven days, on the logic that apples likely are not rotting for much longer than that under a tree and evidenced by the apple rot diameters becoming unmanageably large and leaking after 10 days. We conducted a genome-wide association study (GWAS) using PLINK to identify loci underlying local adaptation at the orchard, variety, and orchard-variety scales.

Reciprocal transplant experiments revealed local adaptation and maladaptation operating at all scales (Subramanian et al., Chapter 3). We define orchard-variety-level (mal)adaptation as the fitness of yeasts on their native orchard and apple variety, relative to their fitness on any other orchard-variety combination. Among the 38 strains that mapped to the *H. uvarum* reference genome, orchard-variety-level (mal)adaptation showed the largest range of log fold-change in performance between native and non-native transplants (LFC range:-0.455 to 0.54, mean ± SE: 0.018 ± 0.033). We also measured orchard-level (mal)adaptation as the relative fitness of yeasts tested on any apple variety from their native orchard, relative to any apple from any other orchard, which ranged from-0.312 to 0.379 (mean ± SE: - 0.003 ± 0.023). Similarly, variety-level (mal)adaptation is the fitness of yeasts on their native apple variety (from any orchard) relative to the alternate apple variety (range:-0.197 to 0.229, mean ± SE: 0.001 ± 0.018) (Fig. 2A,C,E). The smaller variety-level range likely reflects the clonal nature of apple varieties, indicating that strains show limited differentiation across apple varieties.

**Figure 2.**
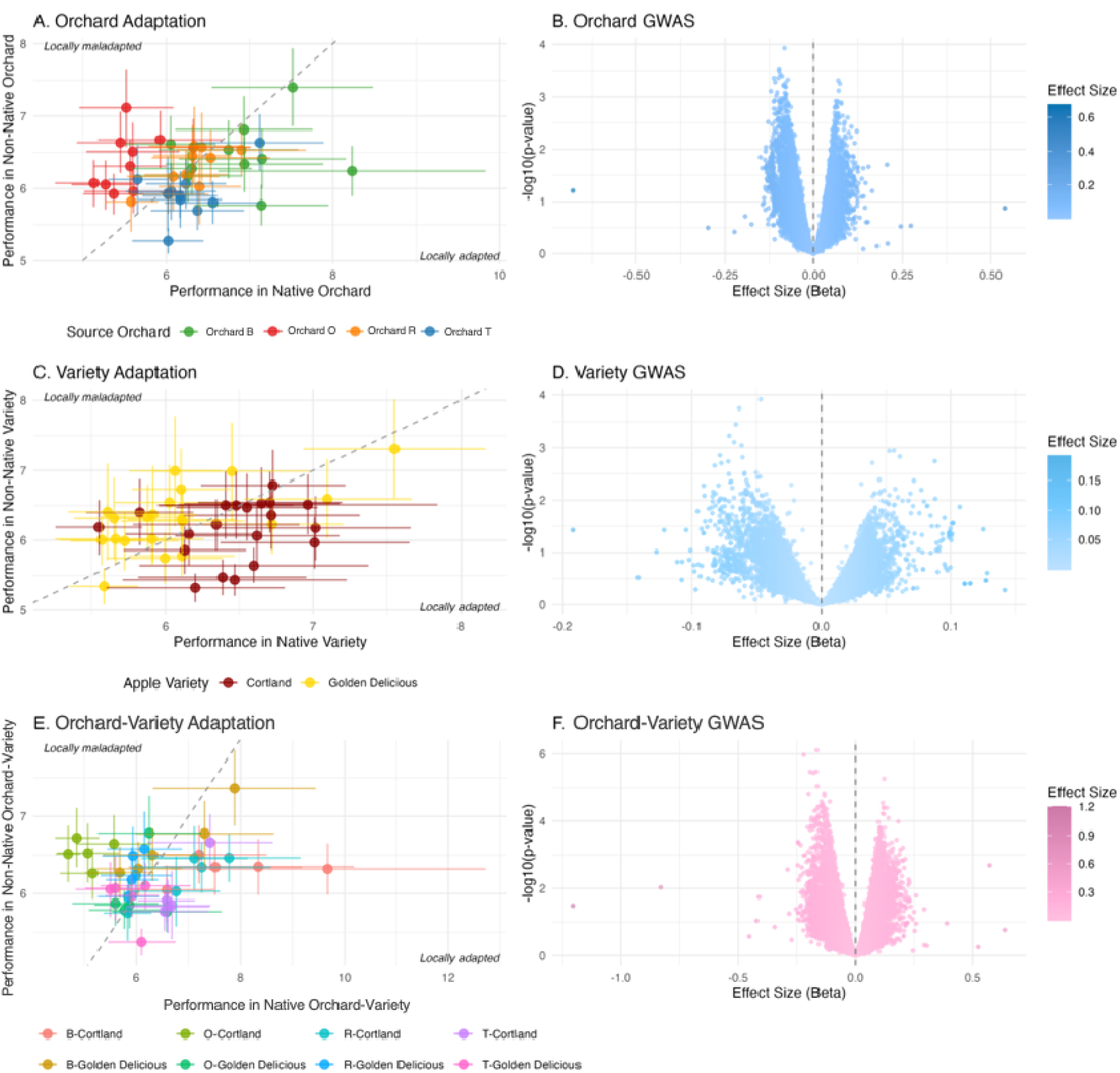
Local adaptation and genome-wide association analysis at multiple ecological scales. (A, C, E) Strain-level adaptation assessment showing performance in native versus non-native environments for orchard (A), variety (C), and orchard-variety combination (E) contexts. Points below the diagonal line indicate local adaptation (better performance in native environment), while points above indicate maladaptation. Strains are colored by source orchard (A), apple variety (B), or orchard-variety combination (C). (B, D, F) Genome-wide association study (GWAS) Manhattan plots showing statistical significance (-log10 p-values) for adaptation and GWAS effect sizes (beta values) for the same orchard (B), variety (D), and orchard-variety combination (F), with the vertical line at zero indicating no effect. Positive beta values indicate alleles conferring better adaptation to the respective environment.

Genome-wide association analysis identified scale-dependent patterns of genetic architecture underlying local adaptation (Fig. 2B,D,F). Orchard-variety adaptation showed the strongest genetic signal with 4,428 significant SNPs (15.1%, *p* < 2e-16 vs. 5% expectation), followed by orchard adaptation with 2,356 significant SNPs (8.0%, *p* < 2e-16), while variety adaptation showed only 675 SNPs (2.3%, *p* = 1.0, not significantly above chance) (SI Appendix, Fig. S3). Effect size (beta) distributions from the GWAS also differed across scales, with orchard-variety adaptation showing the largest range (median β =-0.003, range:-1.202 to 0.635), followed by orchard adaptation (median β =-0.002, range:-0.676 to 0.538), and variety adaptation (median β = 0.001, range:-0.192 to 0.142) (Fig. 2B,D,F; SI Appendix, Fig. S4, top genes in Table S2).

These results suggest scale-dependent patterns of local (mal)adaptation, with strongest (mal)adaptive responses occurring at the orchard-variety level rather than broader orchard or variety scales. The stronger genetic signals for orchard-variety-level adaptation could reflect additive effects of orchard and variety selection or unique selective pressures emerging from non-additive interactions between orchard and variety factors. Rapid, microgeographic adaptation may be occurring, resulting in fitness trade-offs where alleles beneficial in one orchard-variety combination are neutral or detrimental in others, or even detrimental in their own combination due to environmental changes (Brady et al. 2019). Rapid microgeographic adaptation with fitness trade-offs is well-documented in microbial systems (Ferguson et al. 2013). This scale-dependent architecture suggests that environmental context fundamentally shapes both the genetic basis and magnitude of local (mal)adaptation, with finer-scale environmental heterogeneity driving more localized genetic responses despite broader patterns of orchard-variety combination structure.

### Parallel evolution and local adaptation are biased towards high-centrality genes

We tested whether gene network position predicts evolutionary outcomes. By analyzing relationships between network centrality measures from the comprehensive *Saccharomyces cerevisiae* gene-gene interaction network and our genomic evolution/adaptation results. While using the *S. cerevisiae* network data as a proxy for *H. uvarum* gene relationships is a limitation, core genetic interaction patterns are typically conserved across yeast species and eukaryotic organisms, making this a reasonable approximation at this time for identifying broad network effects on evolution (Wuchty et al. 2003, Kellis et al. 2003, Koch et al. 2012, Hahn and Kern 2013). We mapped 2,143 *H. uvarum* genes to *S. cerevisiae* orthologs and calculated four centrality measures: degree (number of direct connections), betweenness centrality (how often a gene lies on paths between other genes), closeness centrality (average distance to all other genes), and eigenvector centrality (connections to highly connected genes). We concatenated metrics of genomic and adaptation from the SNP level to the gene level through methods appropriate for each measure (Methods and Materials).

We found that network centrality systematically affects both evolutionary patterns and adaptive responses. Genes that connect different network modules are more likely to evolve in parallel across apple varieties. This was measured as the correlation between gene-wide average F_ST_ between varieties of apple, versus betweenness centrality (*r* = 0.046, *p* = 0.034) (Fig 3A,C). As a reminder betweenness centrality identifies bow-tie genes that bridge distinct network regions. This positive correlation indicates that genes in connecting positions exhibit more differentiation in response to apple variety selection. This pattern aligns with previous findings that bow-tie position genes evolve in parallel across species, emphasizing the importance of connector genes for repeatable evolution (Friedlander et al. 2015). Despite less overall genomic differentiation between apple varieties, those differences that do occur are concentrated in high-impact, centrally located genes across different orchards, which is supported by other centrality metrics and evolutionary metrics as well (Fig 3A; SI Appendix, Fig. S5).

**Figure 3.**
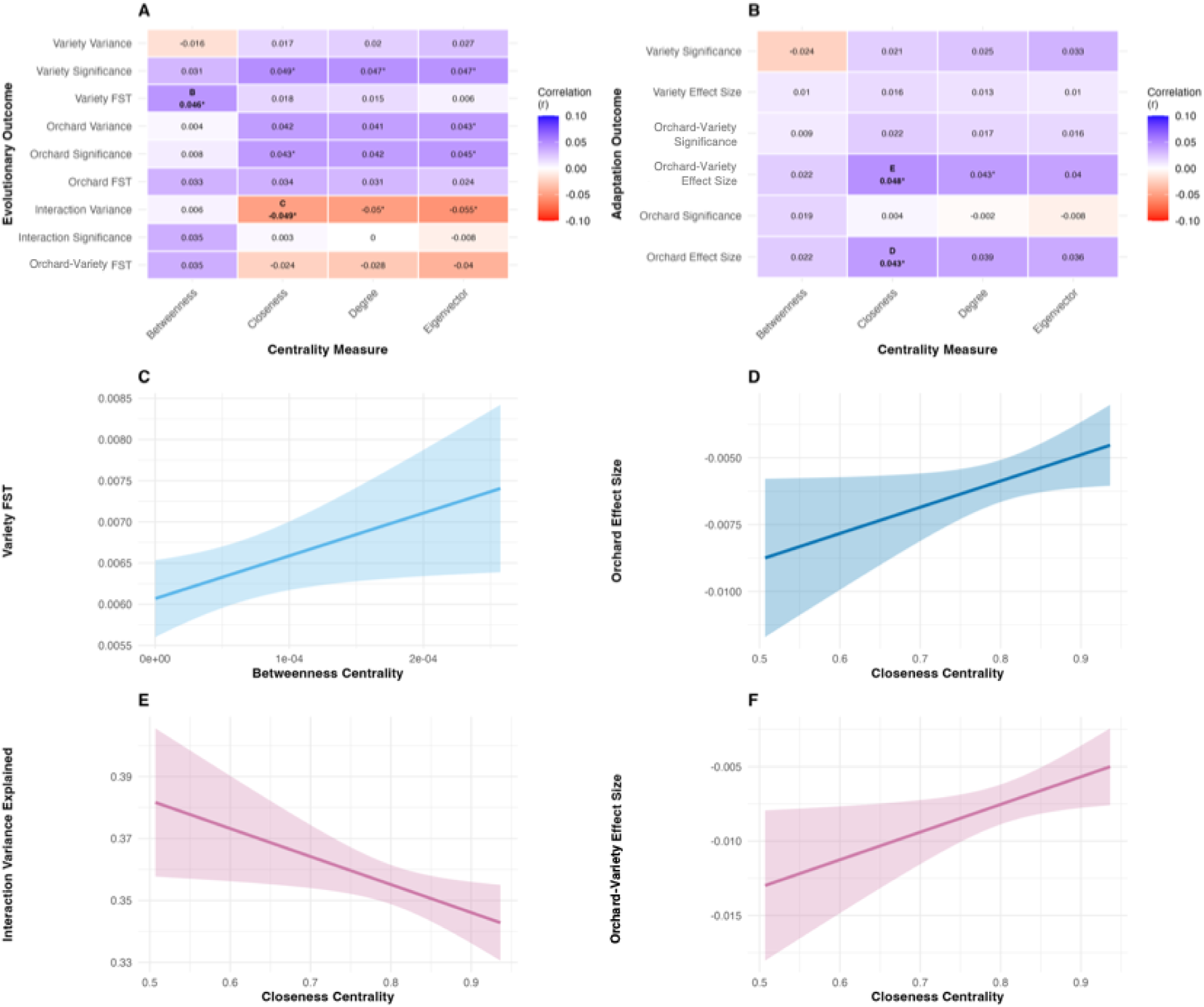
Network centrality affects evolutionary dynamics and local adaptation. (A) Correlation heatmap showing relationships between four centrality measures and nine evolutionary outcomes across *H. uvarum* populations. (C, E) Trendlines of significant evolution-centrality relationships: (C) variety differentiation increases with betweenness centrality, and (E) interaction evolution variance decreases with closeness centrality, points omitted for clarity. (B) Correlation heatmap for adaptation-network relationships showing correlations between centrality measures and GWAS outcomes (significance and effect sizes) for orchard, variety, and orchard-variety adaptation contexts. (D, F) Trendlines of significant adaptation-centrality relationships: orchard (D) and orchard-variety (F) adaptation effect sizes both increase with closeness centrality, points omitted for clarity.

Genes with fewer network connections contribute more to non-parallel evolutionary responses. Non-parallel evolution results in allele frequencies exhibiting an interaction between orchard and apple variety: diverging between varieties in some orchards but not others. Genes with more non-parallel evolution tend to have lower closeness centrality, which measures how central a gene is to the overall network, (*r* =-0.049, *p* = 0.023) (Fig. 3A,E). This suggests that peripheral genes are more likely to be locations of high effect size for orchard-variety-level evolution. This negative correlation held across multiple centrality measures (degree: *r* =-0.050, *p* = 0.021 and eigenvector centrality: *r* =-0.055, *p* = 0.010), indicating that genes that are less connected in the gene-gene interaction network tend to exhibit more idiosyncratic evolution, consistent with theory (citation).

Orchard-level evolution shows an intermediate pattern between variety and orchard-variety responses. Multiple centrality measures correlated positively with orchard evolution signal and effect size on genomic variation (orchard variance with closeness (*r* = 0.042, *p* = 0.050), and eigenvector centrality (*r* = 0.043, *p* = 0.048), and orchard Fisher’s combined p-values with degree (*r* = 0.042, *p* = 0.05), closeness (*r* = 0.043, *p* = 0.045), and eigenvector centrality (*r* = 0.045, *p* = 0.038) (Fig 3A). This suggests that orchard environments provide consistent enough selection pressures for central genes to evolve repeatably, unlike the highly variable orchard-variety-level pressures.

Network position also affects which genes contribute to local adaptation versus maladaptation.

Genes that are more strongly connected are more likely to contribute to local adaptation, rather than maladaptation. At both orchard and orchard-variety levels, GWAS effect sizes showed positive correlations with closeness centrality (orchard adaptation: *r* = 0.043, *p* = 0.047) and orchard-variety adaptation: *r* = 0.048, *p* = 0.027) (Fig. 3B,D,F; SI Appendix, Fig. S6). This suggests that genes occupying central network positions are more likely to harbor alleles with beneficial fitness effects for local conditions, while peripheral genes are more likely to have alleles detrimental to yeast fitness in local conditions.

These patterns demonstrate how epistatic constraints shape evolutionary outcomes in a scale-dependent manner. Genes occupying bow-tie network positions that bridge distinct functional modules facilitate parallel evolution across replicated selective environments (apple varieties across orchards) despite minimal overall genetic differentiation, consistent with theoretical predictions that bow-tie genes are evolutionary hotspots (Stern and Orgogozo 2009, Friedlander et al. 2015). In contrast, peripheral genes with fewer epistatic interactions evolve rapidly to fine-scale environmental heterogeneity but may produce maladaptive outcomes if selection pressures are ephemeral or genotypes are mismatched to environmental change (Alvarez-Ponce et al. 2017, Brady et al. 2019). The intermediate pattern observed for orchard-level evolution, where centrally located genes respond adaptively to consistent selection from orchard management and abiotic factors, suggests that the predictability of evolution depends on both network architecture and the temporal stability of environmental heterogeneity.

### Parallel adaptation to apple variety across orchards: genes, function, and network position

To test whether similar selective pressures result in parallelism at the genetic, functional, and network levels, we identified the top 150 genes with strongest evidence for variety-specific evolution within each orchard and compared these gene sets across orchards. If evolution to apple variety differences is repeatable, we expect genes evolving in response to apple variety within different orchards to be more similar than random gene sets.

Genes evolving in parallel to variety across orchards showed significantly greater similarity than random expectation at multiple biological levels of organization. Gene showed greater genetic similarity compared to random (observed Jaccard similarity = 0.226 vs. random = 0.036, Wilcoxon *p* < 1e-5). This result indicates that the genes evolving in response to variety differences across orchards are far more similar than would be expected of random gene sets, but their similarity is still quite low compared to other measures (Fig. 4). Sets of genes evolving to apple variety also showed greater functional similarity compared to random gene sets (observed GO term Jaccard similarity = 0.761 vs. random = 0.718, *p* < 0.001), suggesting that parallel evolution occurs in genes with more related biological functions even when specific genes differ between orchards (Fig. 4; SI Appendix, Fig. S7). Of the network centrality measures, only betweenness centrality showed significant similarity across the four orchards compared to random gene sets (observed = 0.987 vs. random = 0.976, *p* = 0.042), supporting the hypothesis that genes occupying similar network positions, specifically those in bowtie positions, are somewhat more likely to evolve in parallel.

**Figure 4.**
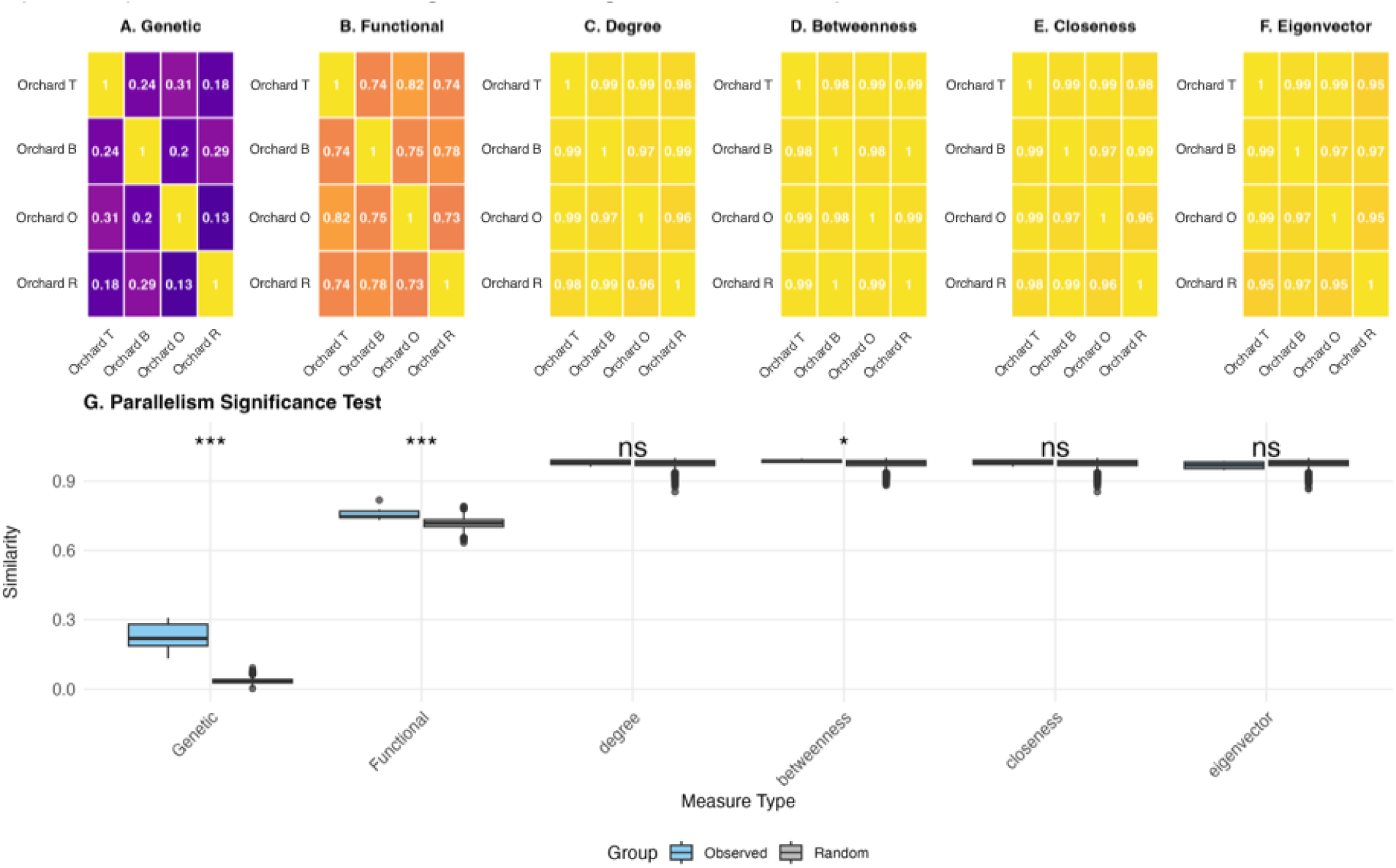
Parallel evolution analysis with similarity matrices. (A-F) Heatmaps showing pairwise similarity between top 150 variety-specific genes identified within each orchard. Similarities calculated using Jaccard index for genetic overlap (A), GO term overlap (B), and cosine similarity of binned centrality distributions for degree (C), betweenness (D), closeness (E), and eigenvector (F). Values range from 0 (no similarity) to 1 (identical). (G) Boxplot comparing observed similarities (blue) to random expectation (gray) from 1,000 permutations. Significance stars indicate results of one-sided Wilcoxon rank-sum tests (*** p < 0.001, ** p < 0.01, * p < 0.05). Random gene sets were generated by sampling genes of equal size from the universe of all genes with network data.

These results suggest that similar selective pressures result in more repeatable evolution than neutral evolution at the genetic, functional, and network levels. The genetic similarity indicates that while some genes evolve repeatedly across orchards in response to variety selection, especially more than random gene sets, most genetic changes are orchard specific (1-Jaccard = 0.774), reflecting the reality that multiple genetic routes can lead to similar adaptive outcomes (Conte et al. 2012, Bolnick et al. 2018). However, the high functional similarity suggests that even when different genes evolve, they tend to be functionally related. This pattern on functional convergence despite genetic divergence is consistent with theoretical predictions that phenotypic evolution is more predictable than genotypic evolution when selection acts on complex traits encoded by redundant genetic pathways (Bolnick et al. 2018).

The greatest similarity across orchards was observed at the network level, specifically for betweenness centrality, supporting the hypothesis that genes occupying bow-tie positions are hotspots for parallel evolution regardless of which specific genes are under selection. This finding provides empirical support for theoretical models predicting that network architecture constrains evolutionary trajectories more strongly than gene-level constraints (Des Marais et al. 2017, Yang and Scarpino 2020). Together, these patterns suggest that epistatic constraints serve as a fundamental organizing principle determining where in the genome parallel evolution is likely to occur. While the genetic targets of parallel selection pressures may vary due to historical contingencies, the functional categories and network positions of evolving genes are far more predictable. This hierarchy of predictability, highest at the network level, intermediate at the functional level, and lowest at the genetic level, indicates that incorporating network architecture into evolutionary forecasting models could substantially improve our ability to predict genomic responses to environmental change, even when we cannot identify which exact genes will evolve.

## Conclusions

Our findings suggest that gene network architecture fundamentally shapes the predictability of evolution in natural orchard-variety combinations, with network centrality determining both the repeatability of evolutionary responses and their adaptive consequences. We show here that highly connected genes occupying bow-tie positions in gene-gene interaction networks are hotspots for parallel evolution across replicated selective environments, while peripheral genes with fewer epistatic interactions facilitate rapid, non-parallel evolutionary responses to fine-scale environmental heterogeneity. While we used the *S. cerevisiae* gene-gene interaction network as a proxy for *H. uvarum* interactions, the deep phylogenetic conservation of core network architectures across yeast species (Wuchty et al. 2003, Kellis et al. 2003, Koch et al. 2012) and eukaryotic organisms (Hahn and Kern 2013) suggests that our findings reflect fundamental evolutionary constraints rather than species-specific idiosyncrasies. Critically, these scale-dependent patterns of network constraint also predict which evolutionary changes contribute to local adaptation versus maladaptation, with central genes responding to consistent selection pressures and harboring alleles with adaptive effects, and peripheral genes evolving rapidly to specific selection pressures and harboring alleles with maladaptive effects. This relationship between network constraints and fitness consequences suggests that while historical contingency and stochasticity remain important, network centrality provides a mechanistic foundation for understanding why evolution follows predictable versus contingent paths in natural populations.

Our results align with accumulating evidence from diverse systems showing that network position predicts evolutionary dynamics. In protein-protein interaction networks, hub proteins evolve slower and show greater sequence conservation (Fraser et al. 2002, Carlson et al. 2006), consistent with our finding that central genes experience stronger evolutionary constraint. Studies of gene regulatory network evolution across yeasts suggest that while specific regulatory interactions can be extensively rewired over evolutionary time, the modular organization and bow-tie architecture of networks remain conserved (Ihmels et al. 2005, Tanay et al. 2005, Tuch et al. 2008, Jovelin and Phillips 2009, Nocedal and Johnson 2015), supporting our finding that network centrality constrains evolutionary paths despite genetic turnover. However, some studies have found weak or context-dependent relationships between connectivity and evolutionary rate (Rogers et al. (manuscript)), suggesting that the network-evolution relationship may depend on the type of network analyzed, the measure of centrality, or the specific selection pressures operating.

Future work integrating network approaches with experimental evolution and comparative genomics across species will be essential for determining whether the network centrality-evolution-adaptation relationship we document in wild yeast represents a general principle of adaptive evolution across organisms, or whether it is contingent on specific features of yeast genetics, microbial life history, or the particular selective environment we studied. The apple orchard system, with its replicated environments and generational timescales, provides a powerful natural experiment for testing these predictions, but it will be important to see if these principles extend broadly across taxa where gene interaction networks can be characterized. These results have important implications for evolutionary forecasting in an era of rapid environmental change. Rather than attempting to predict which specific genes will evolve in response to selection, our network-based framework suggests we can predict where in the genome evolution is most likely to occur and how repeatable those changes will be across populations (Bolnick et al. 2018, Pearless and Freed 2024). For organisms facing anthropogenic stressors, network centrality analyses could identify genetic loci most likely to show parallel responses and adaptive responses, enabling more accurate evolutionary predictions and targeted management strategies. Our findings suggest that incorporating network architecture into evolutionary models could improve predictions of how populations will respond to environmental change.

## Methods and Materials

### Sample Collection and Sequencing

We isolated *Hanseniaspora uvarum* strains from rotting apples collected during fall harvest from four apple orchards across Connecticut: Orchard T, Orchard B, Orchard O, and Orchard R (see coordinates in Subramanian et al. 2025, Chapter 3). We sampled two apple varieties from each orchard: Cortland and Golden Delicious. We focused on *H. uvarum* because community surveys identified it as the most abundant and most variable yeast species across our orchard sites, comprising 4.3% to 97.5% of communities depending on location, making it ideal for studying local adaptation patterns (Subramanian et al. 2025, Chapter 3).

Detailed sample collection protocols are described in Subramanian et al. (2025). We used sterile technique to collect five replicate rotting apples from the ground and fresh apples from a tree for every orchard-variety combination. The five replicates were taken from different trees within each orchard. In the lab, we flame-sterilized a metal inoculating loop and swirled it in the rotting flesh within the punctured rotting apples and streaked the yeast community on a YPD agar plate with 1% penicillin-streptomycin to suppress bacterial growth. After culturing for 24 hours, we isolated individual colonies using sterile pipette tips and cultured them in YPD media. This was followed by BOMB DNA extraction (Oberacker et al. 2019) and species identification using ITS region sequencing with ITS1 and ITS4f primers. From this collection, we selected 40 *H. uvarum* isolates from the five biological replicates per orchard-variety combination.

We extracted high-molecular-weight genomic DNA using the Monarch Genomic DNA Purification Kit (#T3010S). DNA libraries were prepared following PacBio protocols and sequenced on the PacBio Revio system using HiFi technology by the University of Connecticut Center for Genomic Innovation. Sequencing was performed in two batches: high-coverage samples (seven samples, two from Orchard B Golden Delicious apples, two from Orchard O Cortland apples, and three samples from Orchard R Cortland apples, ∼1500× coverage) and standard-coverage samples (33 samples, ∼230-300× coverage) (SI Appendix, Fig S1). We converted raw sequencing data in BAM format to FASTQ using bam2fastq from SMRT Tools (v13.1). We performed quality assessment using FastQC (v0.12.1) and MultiQC (v1.15), with all samples exhibiting >Q20 average base quality scores.

### Population Genomics and Variant Calling

We aligned high-quality PacBio HiFi reads to the *H. uvarum* reference genome (GCA_037102615.1_ASM3710261v1, 9.12 Mb) using minimap2 (v2.24) with parameters optimized for HiFi data. Alignments were sorted and indexed using SAMtools (v1.16.1). Genome-wide coverage analysis confirmed uniform coverage across the 14 major scaffolds, with 38 samples achieving >99% mapping rates to the reference genome (SI Appendix, Fig. S1). Due to low mapping rates, we excluded two samples from Orchard R Cortland apples, resulting in 38 high-quality *H. uvarum* strains for downstream analysis (SI Appendix, Fig. S1).

We called variants using Clair3 (v1.0.4) with haploid-specific parameters optimized for yeast genomes and merged individual sample VCF files using bcftools merge (v1.16). The merged population VCF initially contained 294,454 biallelic SNPs, which we filtered to retain only variants present in at least 2 samples to remove likely sequencing errors while preserving population-specific variation, yielding 236,473 high-quality SNPs for downstream analysis. We then performed linkage disequilibrium pruning using SNPRelate with a correlation threshold of 0.95, to retain maximum allelic diversity while removing redundant variants (Zheng et al. 2012). The final dataset contained 29,489 LD-pruned SNPs for population genomic analyses.

### Population Structure Analysis

We assessed population structure using Principal Component Analysis (PCA) implemented in the adegenet R package on mean-imputed genotype data (Jombart 2008). We then performed Discriminant Analysis of Principal Components (DAPC) to identify genetic variation that best separates predefined groups (orchards and apple varieties). For each grouping variable, we used cross-validation with 100 repetitions to determine the optimal number of principal components to retain before discriminant analysis. We tested the statistical significance of genetic differentiation using PERMANOVA with 999 permutations and Euclidean distance on the genotype matrix (Okansen et al. 2022).

We quantified orchard-variety differentiation using F_ST_ calculated with the basic.stats function from the hierfstat R package at three hierarchical levels: among all 8 orchard-variety combinations, among orchards, and between apple varieties (Goudet and Jombart 2022). We analyzed genomic differentiation patterns using binomial generalized linear models for each SNP, testing effects of orchard, variety, and their interaction as fixed effects. We also calculated proportional variance explained for each SNP to get an effect size of orchard, variety, and interaction on allele frequency, using the ratio of factor-specific chi-square statistics over total model deviance from the binomial GLM.

### Local Adaptation Experiments and GWAS

We quantified local adaptation through reciprocal transplant experiments testing all 38 sequenced *H. uvarum* strains across all 8 orchard-variety combinations, following protocols detailed in Subramanian et al. 2025 (Chapter 3). We sterilized fresh apples with ethanol, created standardized inoculation holes using sterile pipette tips, and inoculated with OD600-normalized yeast cultures. After sealing holes with microporous film, we measured rot diameter after 7 days as a measure of yeast performance and fitness. We then calculated local adaptation phenotypes as log fold-change in performance between native and non-native transplants across orchard, variety, and orchard-variety combination.

We performed genome-wide association studies (GWAS) using PLINK (v1.9) with the LD-pruned SNP dataset. We controlled for population structure using the first principal component as a covariate in linear regression models implemented in PLINK v1.9. We tested additional principal components and kinship matrix-based mixed models, but these approaches either failed to converge or provided no improvement in model fit given our sample size and population structure. We used linear regression with additive genetic effects for each adaptation phenotype.

### Network Centrality Analysis

We obtained gene network centrality data from the comprehensive *Saccharomyces cerevisiae* genetic interaction network (Costanzo et al. 2016), which quantifies epistatic relationships among ∼6,000 genes based on systematic double-mutant fitness measurements. We constructed the yeast genetic interaction network from four publicly available datasets from the Costanzo et al. (2016) (SGA_ExE, SGA_NxN, SGA_ExN_NxE, and SGA_DAmP) using the igraph R package (Csardi and Nepusz 2006). We combined 18.9 million pairwise genetic interactions into a single undirected network after removing multiple edges and self-loops. For genes with both temperature-sensitive (ts) and DAmP alleles, only DAmP alleles were retained. We used this network and igraph functions to calculate four unweighted centrality measures for each gene: degree, betweenness centrality, closeness centrality, and eigenvector centrality.

We mapped *H. uvarum* genes to *S. cerevisiae* orthologs using a two-step approach. First, *H. uvarum* proteins identified through BLAST annotation were matched to RefSeq protein identifiers from the *S. cerevisiae* reference proteome (GCF_000146045.2), then linked to systematic gene names and common gene names using the Saccharomyces Genome Database cross-reference table. This mapping yielded network centrality data for 2,143 *H. uvarum* genes with clear *S. cerevisiae* orthologs.

We then assigned SNPs to genes based on physical position within gene boundaries and calculated gene-level statistics. We aggregated p-values of multiple SNPs within a gene from the binomial GLM and the adaptation GWAS using Fisher’s combined probability method to get gene-level p-values. We took the arithmetic means of the proportional variance explained by each factor across SNPs within a gene for effect sizes. We calculated F_ST_ calculated at the gene level as the mean F_ST_ across all SNPs within gene boundaries across orchard, variety, and orchard-variety levels. We took the arithmetic mean of beta across SNPs within a gene from the adaptation GWAS. We evaluated correlations between network centrality measures and evolutionary/adaptation measures using Pearson correlation tests (cor.test function) to obtain correlation coefficients and linear regression models (lm function) to obtain statistical significance (p-values).

### Parallel Evolution Across Multiple Levels of Organization Analysis

We identified gene sets evolving under similar selective pressures by analyzing variety-specific evolution separately within each of the four orchards. For each orchard, we ran binomial GLMs testing SNP associations with apple variety, calculated gene-level significance using Fisher’s combined probability method, and selected the top 150 genes by gene-level p-value as the representative genes for variety specific evolution within that orchard context. We chose 150 genes per orchard based on testing gene set sizes from 50-300 genes and found that smaller sets yielded overly permissive significance levels, while larger sets showed no significant similarity compared to random sets. The 150 gene cutoff provided detectable but stringent significance levels.

We quantified similarity between gene sets using three measures: genetic similarity (Jaccard coefficient of gene overlap), functional similarity (Jaccard coefficient of overlapping GO terms), and network centrality similarity (cosine similarity of quintile binned centrality distributions). We assessed statistical significance through permutation testing with 1000 random gene sets of equivalent size (150 genes each) sampled from the all genes with network centrality data. We then conducted Wilcoxon rank-sum tests to compare observed similarities to random expectations.

## Acknowledgements

We thank the orchard owners and managers for access to sampling sites and allowing us to collect rotting and fresh apples. This research was supported by the National Science Foundation Graduate Research Fellowship, the EEB Graduate Research Fund to the Department of Ecology and Evolutionary Biology and Connecticut State Museum of Natural History, and the NSF Rules of Life Emerging Networks grant (FAIN-2133740). We are grateful to all Bolnick lab members for feedback on experimental design and support.. We would also like to thank Vikas Sarathy for providing assistance during apple picking. We are also grateful to Bo Reese and the University of Connecticut Center for Genomic Innovation for support with genomic sequencing and troubleshooting.

## Supplemental Figures and Tables

**Supplemental Figure S1:**
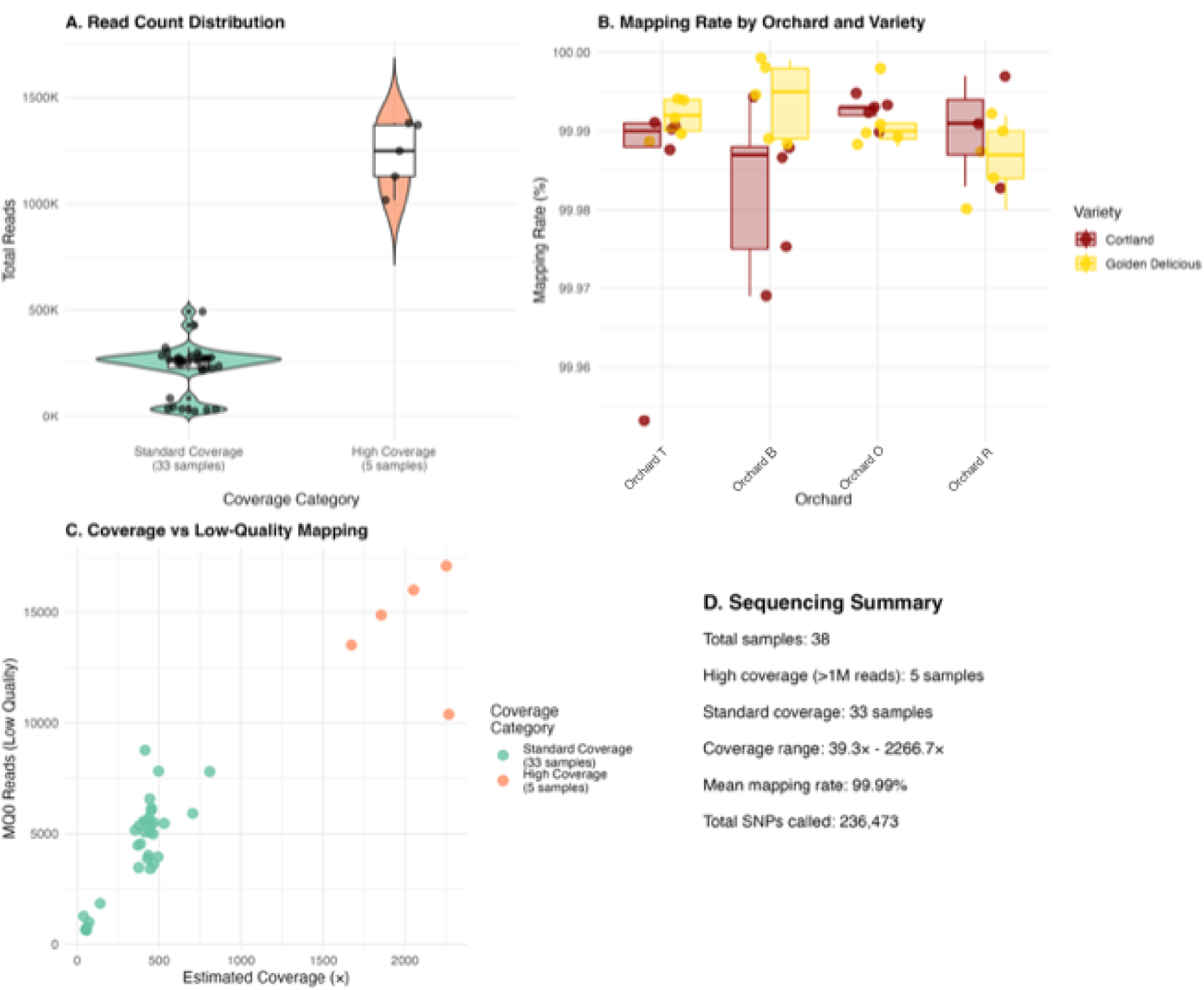
Sequencing Quality Control Metrics. (A) Violin plots showing distribution of total reads for standard coverage (33 samples) and high coverage (5 samples) groups. Individual samples shown as points. (B) Mapping rates across orchards and varieties, with points representing individual samples colored by variety. (C) Estimated genome coverage versus low-quality (MQ0) reads, colored by coverage category. Coverage estimated assuming 15 kb average read length and 9.13 Mb genome size. (D) Summary statistics panel showing key sequencing metrics including total samples, coverage distribution, mean mapping rate, and total SNPs identified. Two samples (RC.1.7 and RC.2.3) were excluded from analysis due to low mapping rates (<30%).

**Supplemental Figure S2:**
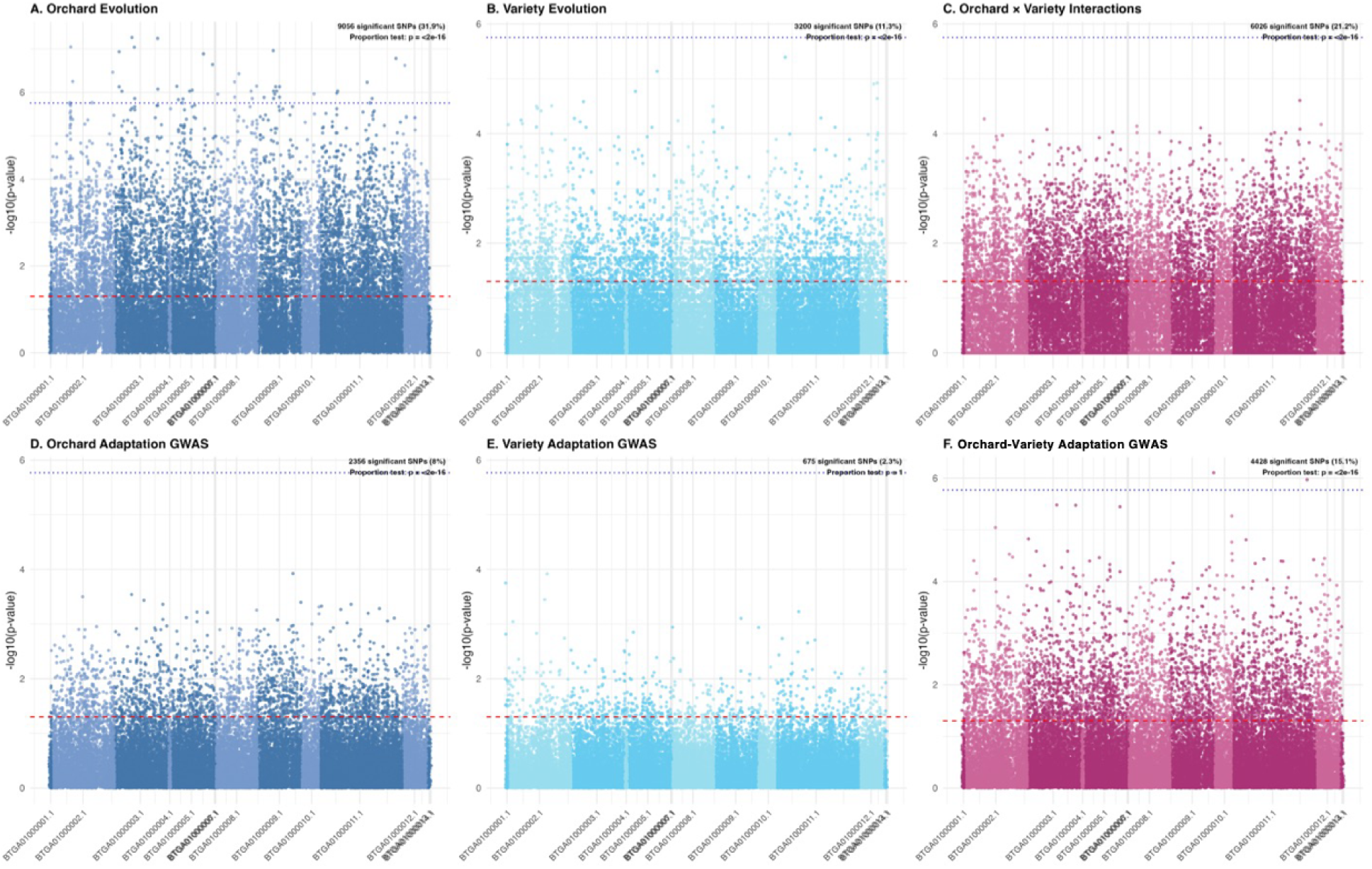
Manhattan Plots with Significance Testing. (A-C) Binomial GLM results for orchard evolution (A), variety evolution (B), and orchard×variety interactions (C). Red dashed line indicates nominal p = 0.05; blue dotted line indicates Bonferroni-corrected threshold. Annotations show number of significant SNPs and results of proportion tests comparing observed significance rate to 5% null expectation. (D-F) GWAS results for orchard adaptation (D), variety adaptation (E), and orchard-variety adaptation (F) with same significance thresholds. Proportion tests assess whether observed rates of significant associations exceed expectations under the null hypothesis of no true associations.

**Supplemental Figure S3:**
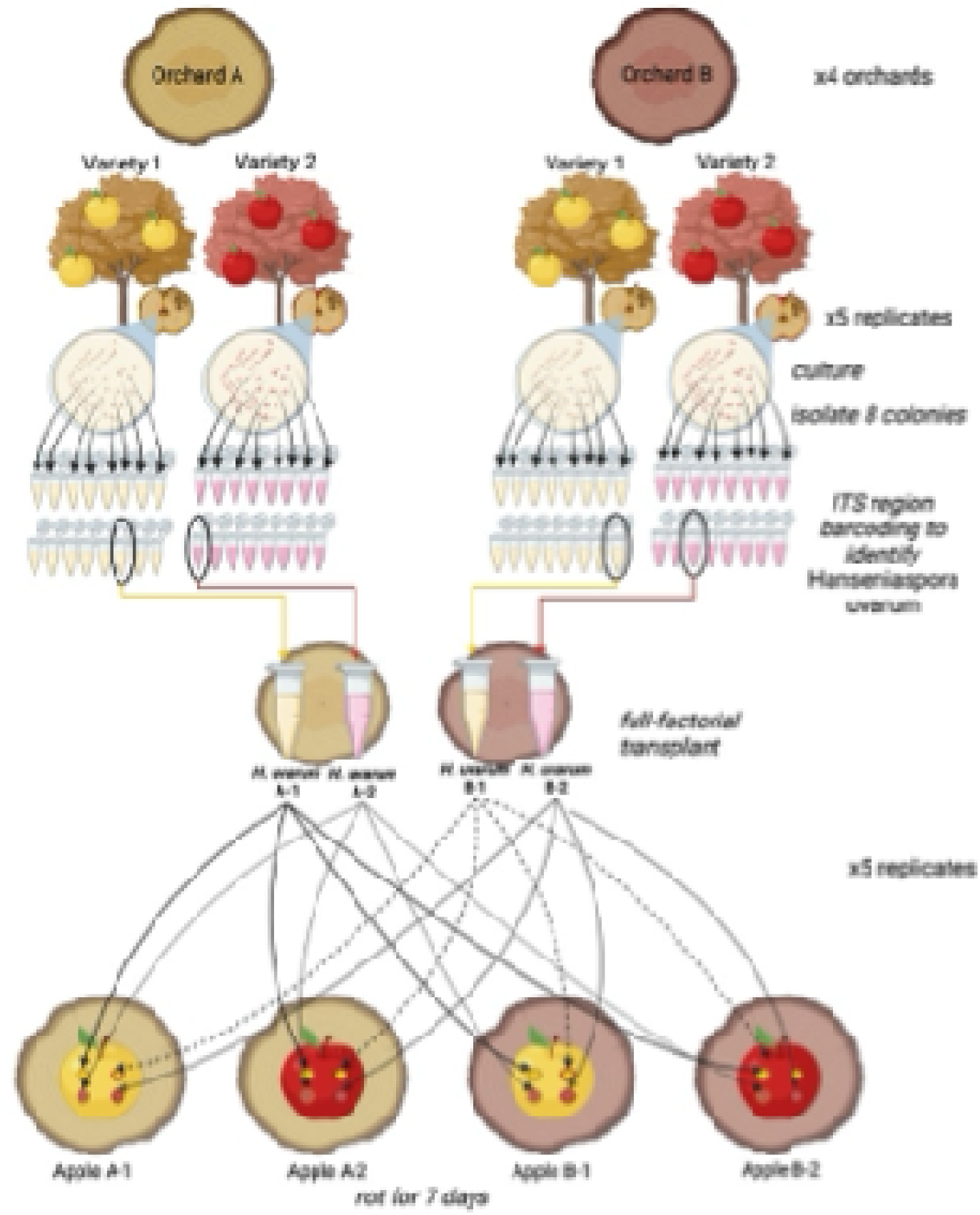
Local adaptation reciprocal transplant experimental method. Created in BioRender.

**Supplementary Figure S4:**
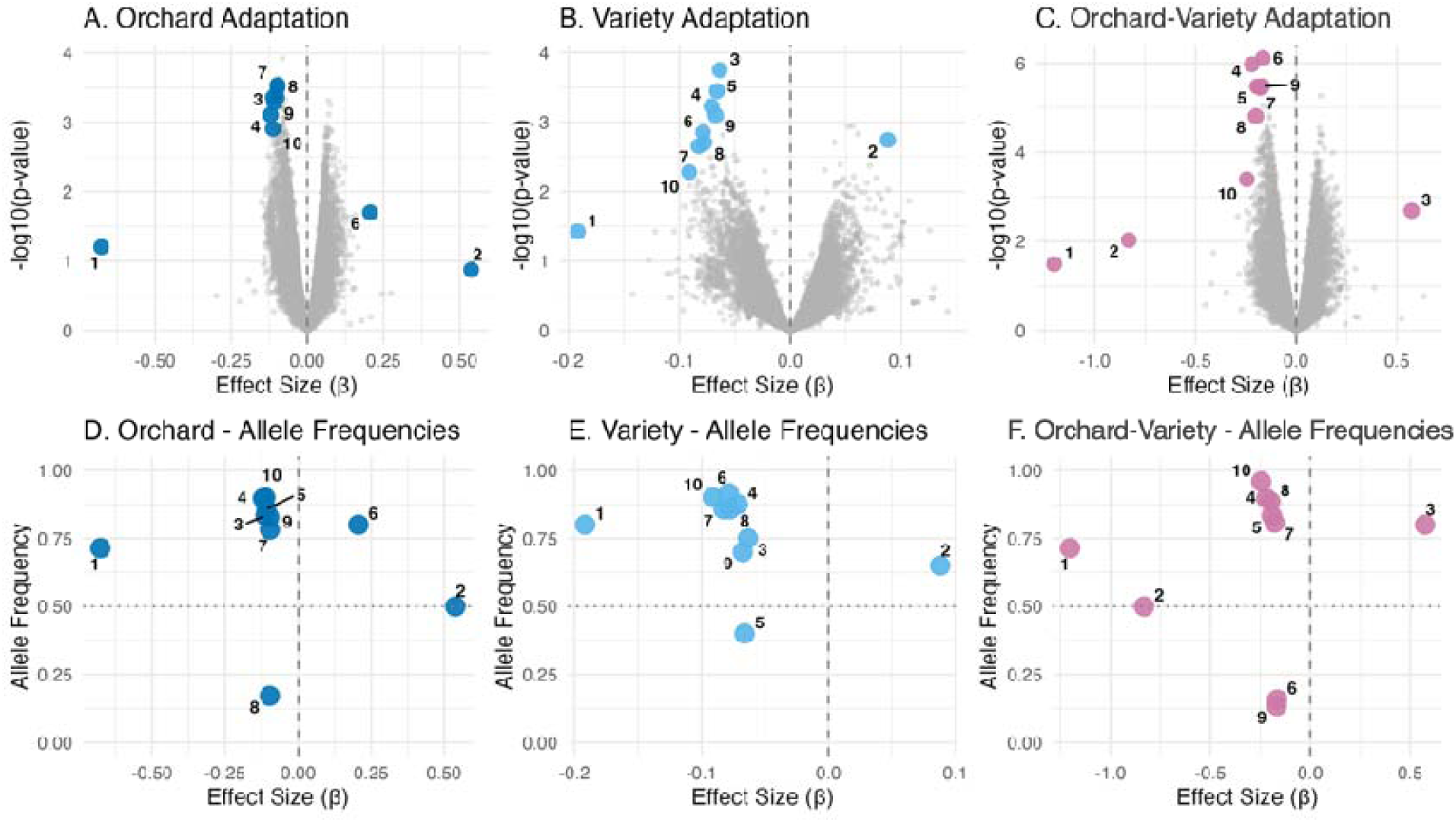
Extreme GWAS Effects and Allele Frequencies. Top panels (A-C) show volcano plots with the top 10 SNPs (by effect magnitude = |β| ×-log10(p)) highlighted for orchard (A), variety (B), and orchard-variety (C) adaptation. Bottom panels (D-F) show allele frequencies of these extreme SNPs versus their effect sizes. Vertical dashed line at β = 0 separates beneficial (positive) from deleterious (negative) effects. Horizontal dotted line at 0.5 indicates intermediate frequency. Numbers correspond to SNPs ranked by effect magnitude.

**Supplemental Figure S5:**
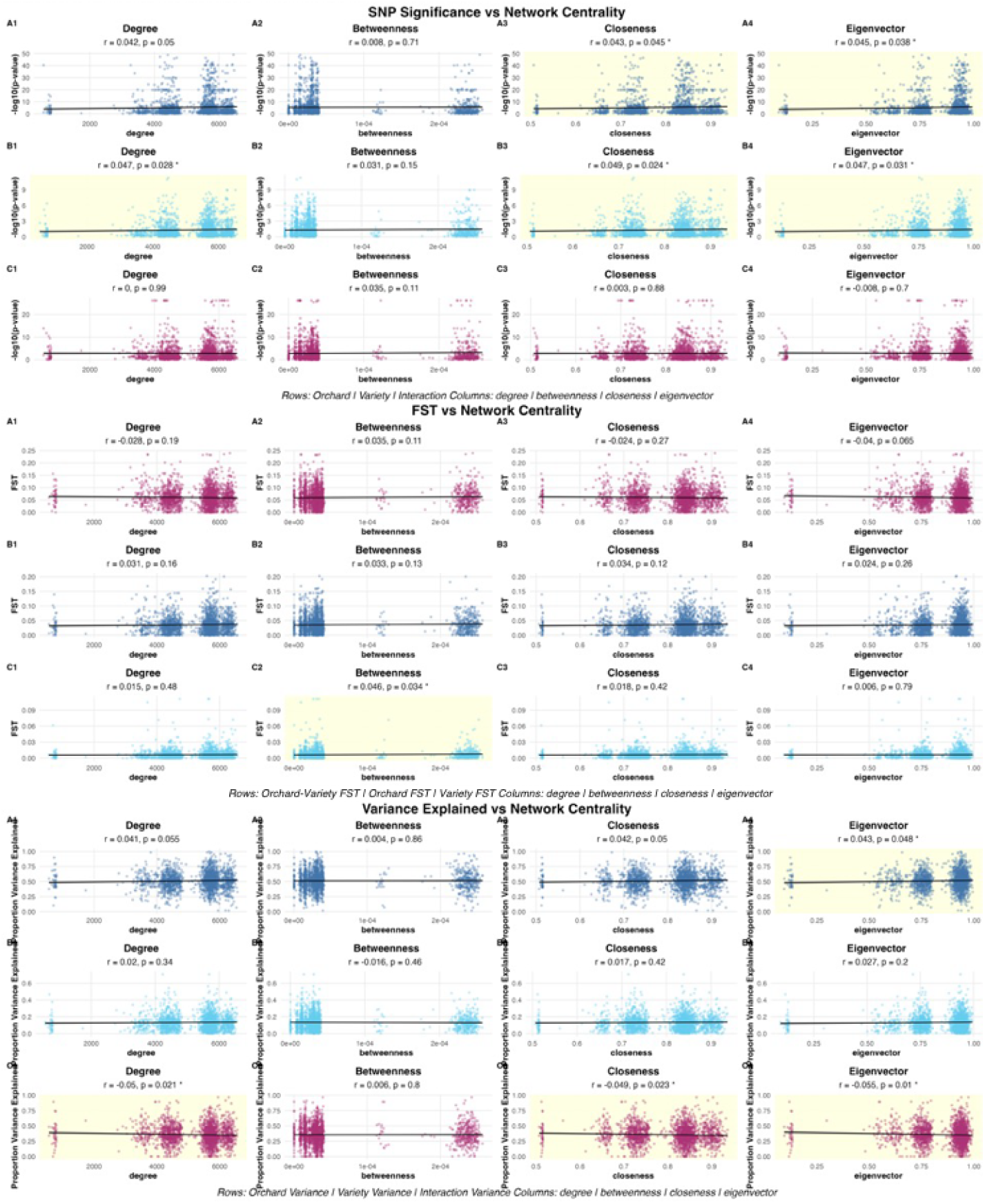
Evolution-Centrality Correlation Scatterplots. Scatterplots show correlations between four centrality measures (degree, betweenness, closeness, eigenvector; columns) and three evolutionary outcomes (rows). Top section: Fisher’s combined p-values aggregated to gene level from binomial GLM results. Middle section: FST calculated across 8 orchard-variety combinations, by orchard, and by variety. Bottom section: Proportion of variance explained by orchard, variety, and their interaction from binomial GLM. Each panel shows individual genes as points with linear regression line and 95% confidence interval. Significant correlations (p < 0.05) highlighted with yellow background.

**Supplemental Figure S6:**
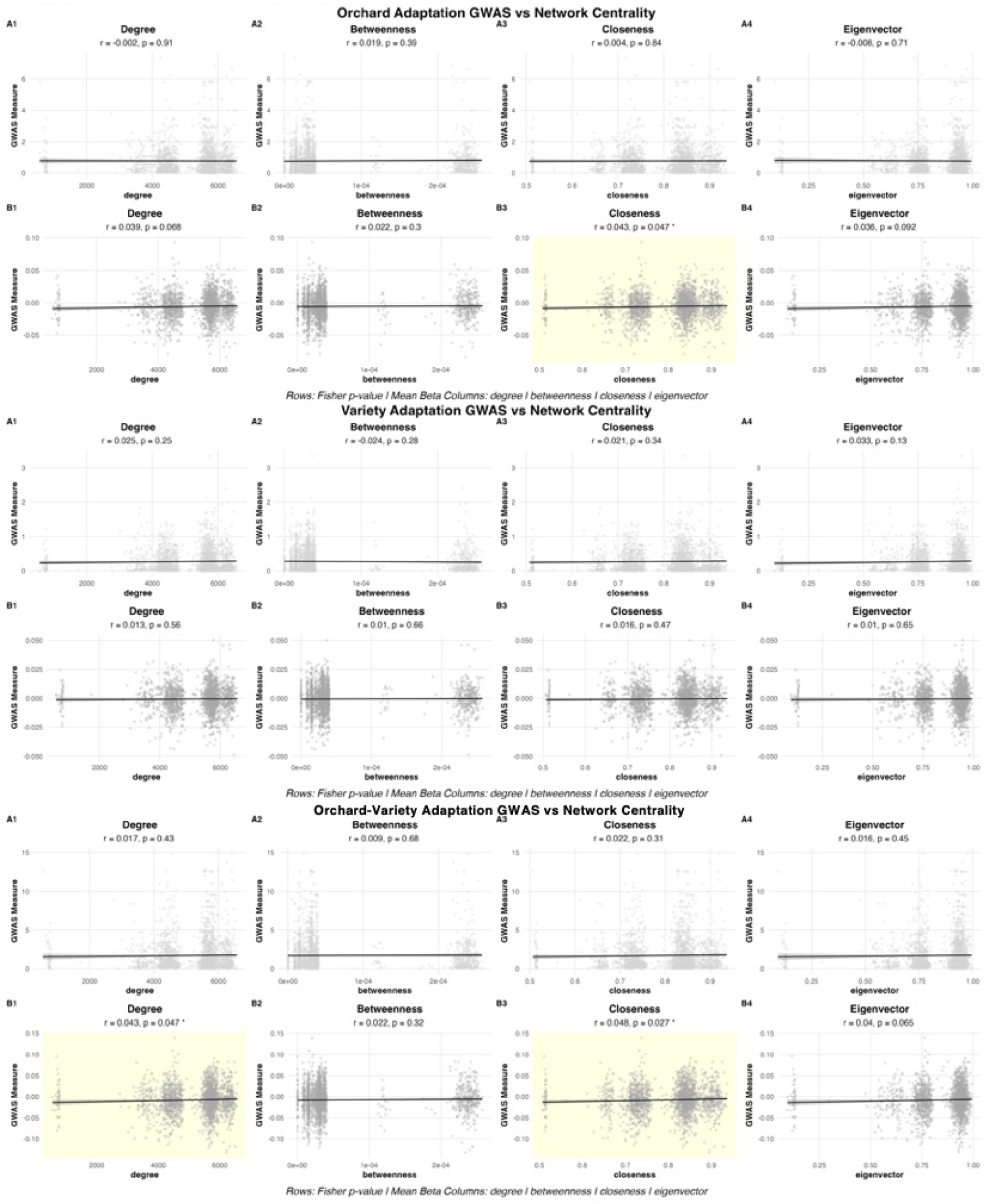
Adaptation-Centrality Correlation Scatterplots. Three sections showing correlations between centrality measures and GWAS results for orchard adaptation (top), variety adaptation (middle), and orchard-variety adaptation (bottom). For each adaptation type, two rows show correlations with Fisher’s combined p-value and mean effect size (β). Layout identical to Supplemental Figure S5, with 8 panels per adaptation type (4 centrality measures × 2 GWAS outcomes). Significant correlations highlighted with yellow background.

**Supplemental Figure S7:**
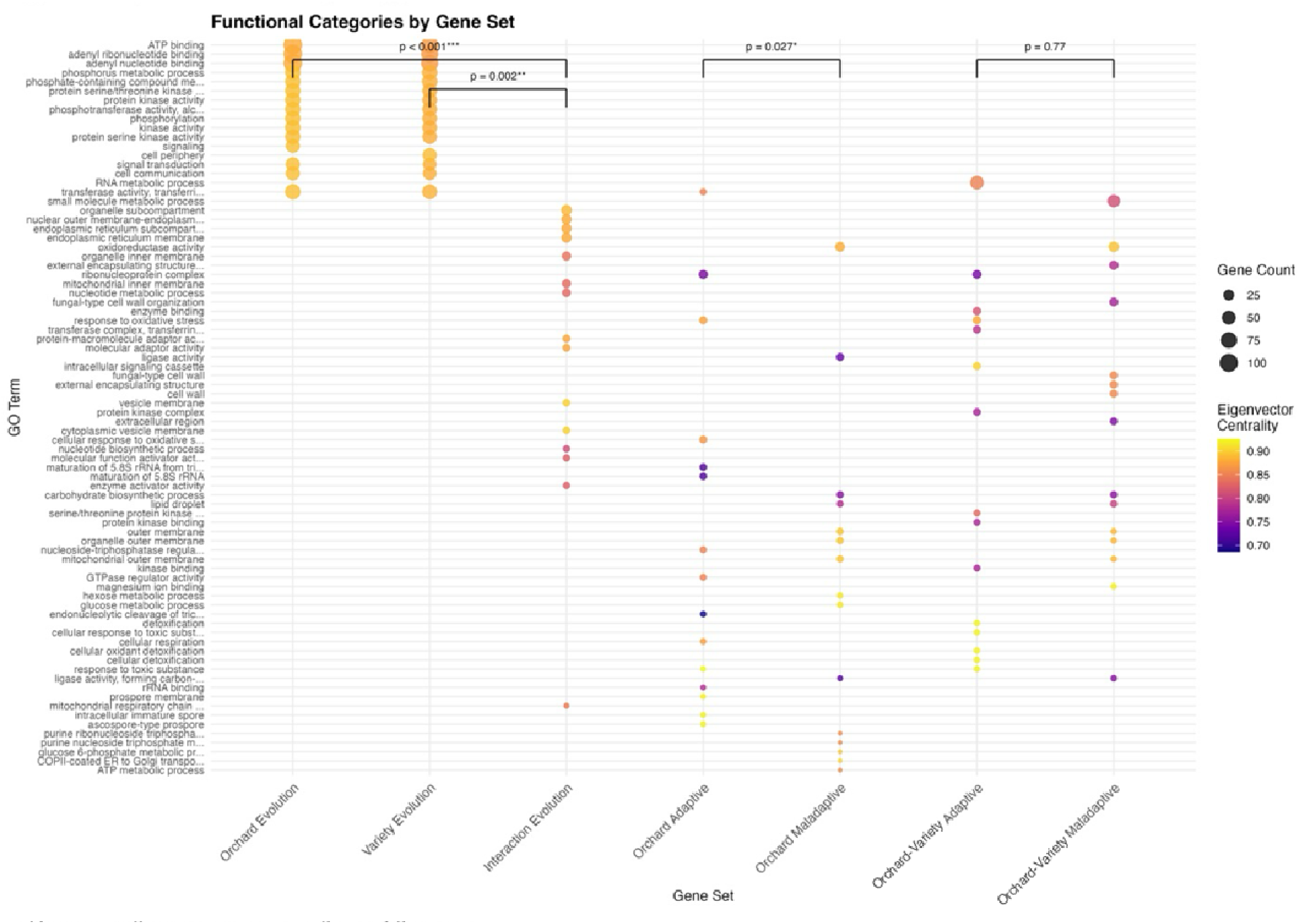
Functional Categories by Gene Set. Dot plot showing top 10 GO terms (by gene count) for each gene set. Y-axis shows GO term descriptions (shortened to 35 characters); x-axis shows seven gene sets ordered as: Orchard Evolution, Variety Evolution, Interaction Evolution, Orchard Adaptive, Orchard Maladaptive, Orchard-variety Adaptive, Orchard-variety Maladaptive. Point size indicates number of genes in the term; point color indicates mean eigenvector centrality of genes in that term. Brackets above indicate significant differences in eigenvector centrality between gene sets based on Wilcoxon tests, with p-values and significance stars shown. Gene sets defined as top 150 genes by Fisher’s combined p-value for evolutionary analyses and by effect size for adaptation analyses.

---

**Supplemental Table S1.** Top genes associated with evolution at multiple ecological scales. Table showing top 10 genes (by Fisher’s combined p-value) for orchard evolution, variety evolution, and interaction evolution. Columns include systematic gene name, gene ID, evolution factor (orchard, variety, and interaction) Fisher’s combined p-value (by factor), and four centrality measures (degree, betweenness, closeness, eigenvector) from the *S. cerevisiae* genetic interaction network.

| Systematic Gene Name | Gene ID | Evolution Factor | Fisher's combined p-value | degree | betweenness | closeness | eigenvector |
| --- | --- | --- | --- | --- | --- | --- | --- |
| YAR019C | NP_009411.2 | Orchard | <0.001 | 6451 | 2.50E-04 | 0.922 | 0.964 |
| YDR507C | NP_010795.3 | Orchard | <0.001 | 4417 | 2.00E-05 | 0.728 | 0.760 |
| YPL153C | NP_015172.1 | Orchard | <0.001 | 4179 | 1.00E-05 | 0.711 | 0.720 |
| YCL024W | NP_009907.2 | Orchard | <0.001 | 4691 | 2.00E-05 | 0.749 | 0.790 |
| YDR490C | NP_010778.3 | Orchard | <0.001 | 5591 | 3.00E-05 | 0.829 | 0.925 |
| YNL298W | NP_014101.1 | Orchard | <0.001 | 4613 | 2.00E-05 | 0.743 | 0.793 |
| YOL113W | NP_014528.1 | Orchard | <0.001 | 5658 | 3.00E-05 | 0.835 | 0.935 |
| YPL140C | NP_015185.1 | Orchard | <0.001 | 5626 | 3.00E-05 | 0.832 | 0.929 |
| YDR477W | NP_010765.3 | Orchard | <0.001 | 4321 | 1.00E-05 | 0.721 | 0.744 |
| YKL166C | NP_012755.1 | Orchard | <0.001 | 5809 | 3.00E-05 | 0.851 | 0.957 |
| YGR233C | NP_011749.3 | Variety | <0.001 | 4474 | 2.00E-05 | 0.733 | 0.754 |
| YCR072C | NP_009997.2 | Variety | <0.001 | 4442 | 2.00E-05 | 0.730 | 0.765 |
| YMR229C | NP_013956.1 | Variety | <0.001 | 6262 | 2.40E-04 | 0.900 | 0.934 |
| YKL213C | NP_012709.1 | Variety | <0.001 | 6521 | 2.50E-04 | 0.931 | 0.987 |
| YOR336W | NP_014981.1 | Variety | <0.001 | 6276 | 2.40E-04 | 0.901 | 0.938 |
| YER078C | NP_011001.1 | Variety | <0.001 | 5713 | 3.00E-05 | 0.841 | 0.946 |
| YPL140C | NP_015185.1 | Variety | <0.001 | 5626 | 3.00E-05 | 0.832 | 0.929 |
| YOR231W | NP_014874.1 | Variety | <0.001 | 5763 | 4.00E-05 | 0.846 | 0.944 |
| YCL024W | NP_009907.2 | Variety | <0.001 | 4691 | 2.00E-05 | 0.749 | 0.790 |
| YDR507C | NP_010795.3 | Variety | <0.001 | 4417 | 2.00E-05 | 0.728 | 0.760 |
| YBR237W | NP_009796.1 | Interaction | <0.001 | 6179 | 2.40E-04 | 0.890 | 0.922 |
| YDL084W | NP_010199.1 | Interaction | <0.001 | 6094 | 2.30E-04 | 0.881 | 0.911 |
| YDL031W | NP_010253.2 | Interaction | <0.001 | 4062 | 1.00E-05 | 0.702 | 0.701 |
| YDR021W | NP_010304.3 | Interaction | <0.001 | 6026 | 2.20E-04 | 0.873 | 0.901 |
| YDR194C | NP_010480.1 | Interaction | <0.001 | 4014 | 1.00E-05 | 0.699 | 0.694 |
| YDR243C | NP_010529.3 | Interaction | <0.001 | 6111 | 2.20E-04 | 0.883 | 0.917 |
| YGL171W | NP_011344.1 | Interaction | <0.001 | 4369 | 2.00E-05 | 0.725 | 0.751 |
| YHR065C | NP_011932.2 | Interaction | <0.001 | 3941 | 1.20E-04 | 0.694 | 0.547 |
| YJL138C | NP_012397.1 | Interaction | <0.001 | 5807 | 4.00E-05 | 0.850 | 0.950 |
| YJL033W | NP_012501.1 | Interaction | <0.001 | 4313 | 2.00E-05 | 0.720 | 0.741 |

**Supplemental Table S2.**
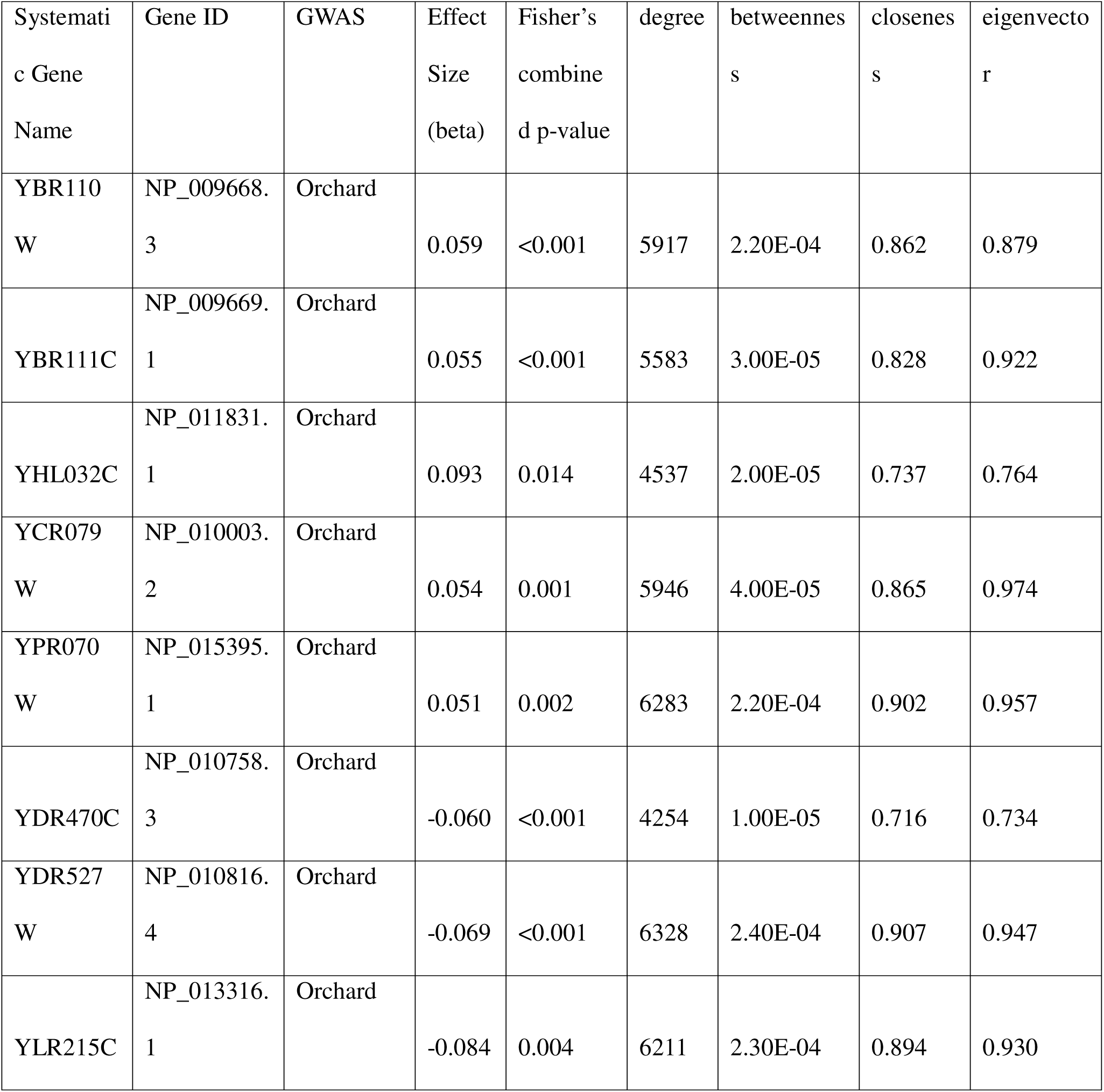
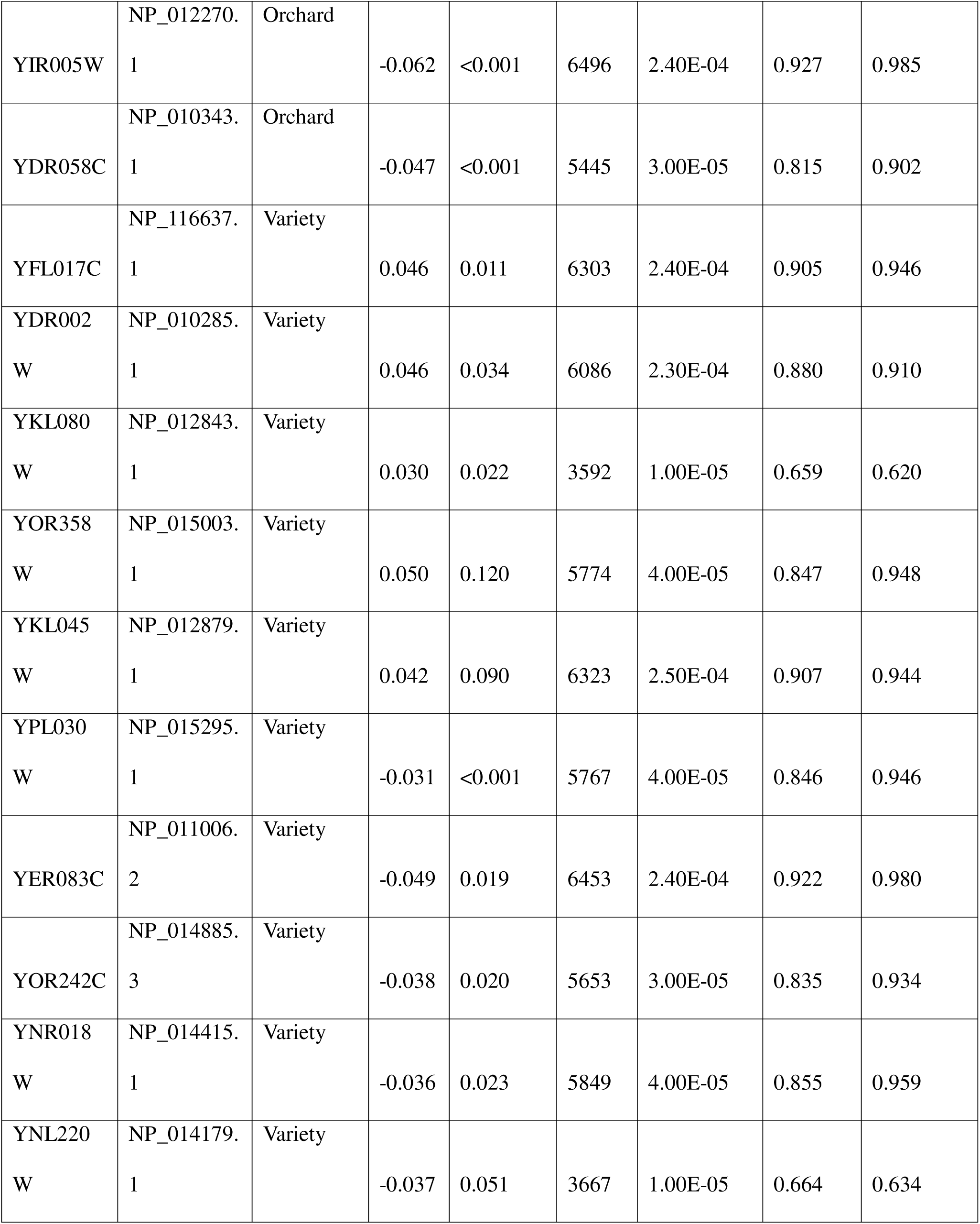
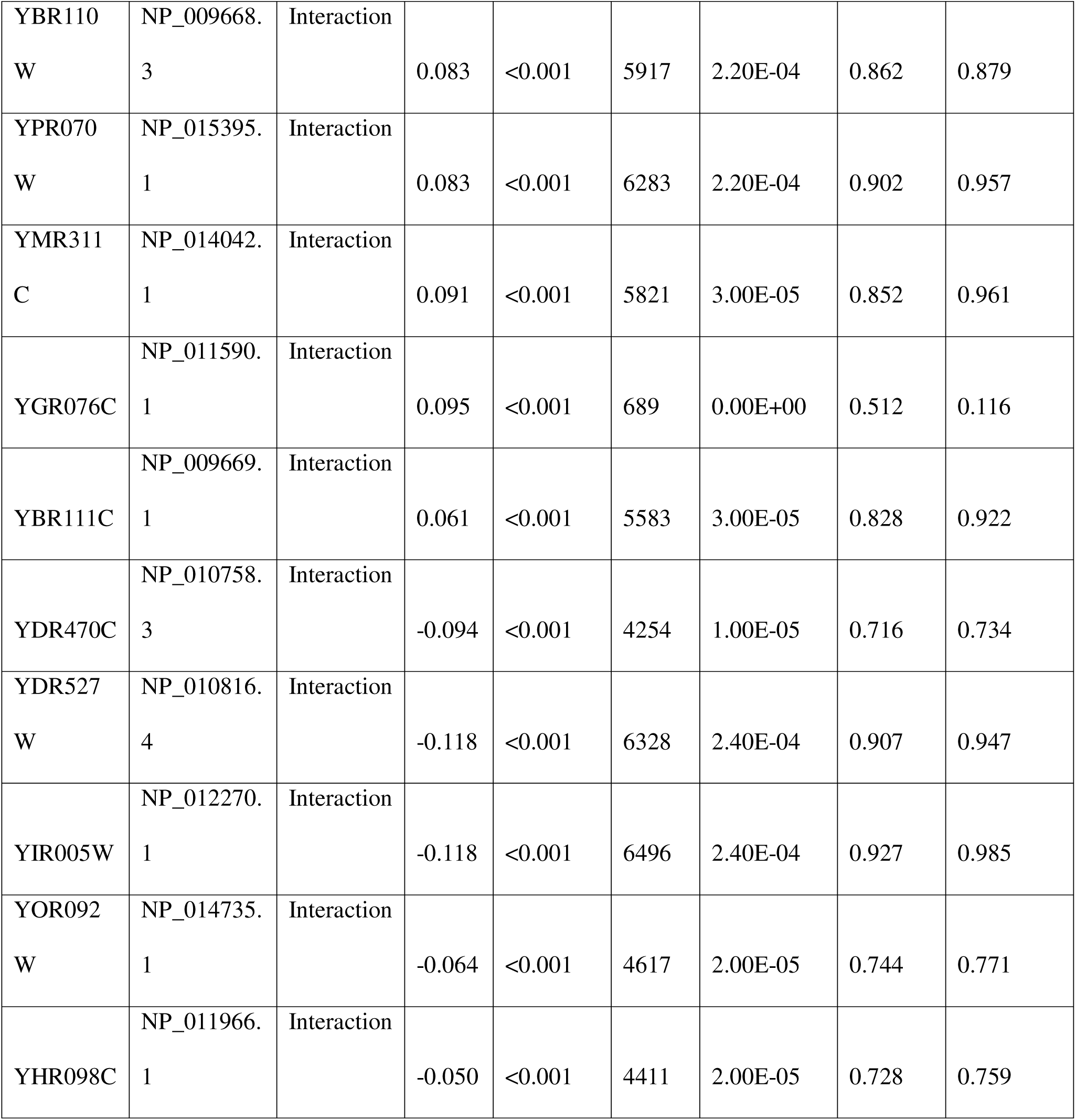
Top genes associated with local adaptation ranked by effect magnitude. Table showing top 5 adaptive and top 5 maladaptive genes (ranked by effect magnitude = |β| ×-log10(p)) for orchard, variety, and orchard-variety adaptation GWAS. Columns include systematic gene name, gene ID, mean effect size (β), Fisher’s combined p-value, number of SNPs per gene, and four centrality measures (degree, betweenness, closeness, eigenvector) from the *S. cerevisiae* genetic interaction network.

## Notes

### Competing Interest Statement

The authors have declared no competing interest.

